# Lipid droplets promote the aberrant liquid-liquid phase separation of alpha-synuclein leading to impaired energy homeostasis

**DOI:** 10.1101/2025.11.20.689615

**Authors:** Jose Cevallos, Elena Eubanks, Sunghoo Jung, Yiming Huang, Elyse Guadagno, Neeharika Rao Suvvari, Aryan Doshi, Nitya Ravinutala, Alejandro Mosera, Nagendran Ramalingam, Arati Tripathi, Tim Bartels, Ulf Dettmer, Eleanna Kara

**Author notes:** Equal contribution (Jose Cevallos, Elena Eubanks). Equal contribution (Sunghoo Jung, Yiming Huang, Elyse Guadagno).

## Abstract

Alpha-synuclein (αSyn) inclusions, termed Lewy bodies, are the characteristic neuropathological feature of Parkinson’s disease. Growing evidence points towards a role of aberrant liquid-liquid phase separation in the dysregulation of αSyn and sequence of events that lead to the formation of Lewy bodies. However, the triggers leading to aberrant phase separation are unknown, as is the relevance of this phenomenon to the neurodegeneration process. In this study, we showed that αSyn spontaneously phase separates into condensates in the presence of lipid droplets. These lipid droplet-rich condensates represent a toxic species of αSyn that prevents the turnover of the entrapped lipid droplets; they are also toxic to neighbouring mitochondria which are depolarized and undergo increased mitophagy. These findings underscore the increasing importance of lipid droplets in the pathogenesis of neurodegenerative diseases, and Parkinson’s disease in particular. The lipid droplets are significantly enriched within the neuromelanin in midbrain dopaminergic neurons in the substantia nigra and could therefore uniquely facilitate the early αSyn-associated neurodegeneration of this region in PD. Our findings reveal a novel pathway implicated in the dysregulation of αSyn that connects aberrant liquid-liquid phase separation, lipid droplets and mitochondrial toxicity.

## Introduction

It is thought that aberrant liquid-liquid phase separation (LLPS) is the first step in the process leading to the formation of proteinaceous inclusions that are involved in the pathogenesis of neurodegenerative diseases. The amyloidogenic proteins involved in these diseases either independently phase separate or partition (get recruited) into a pre-existing condensate that is formed by other proteins (Visser *et al*, 2024). These condensates then undergo a liquid-to-solid transition because of locally increased protein concentration leading to aggregation (Dai *et al*, 2024; Putnam *et al*, 2023). This process eventually leads to the formation of the characteristic proteinaceous inclusions seen in neurodegenerative diseases. This phenomenon has been well studied for tau (Boyko *et al*, 2019; Hochmair *et al*, 2022; Wegmann *et al*, 2018) and TAR DNA-binding protein 43 (TDP-43) (Schmidt & Rohatgi, 2016; Yan *et al*, 2024).

There is recent evidence suggesting that aberrant LLPS of alpha-synuclein (αSyn) is involved in its dysregulation in Parkinson’s disease (PD) and Dementia with Lewy bodies (DLB), eventually leading to the formation of Lewy bodies. αSyn is an intrinsically disordered protein that contains two low complexity domains in its N-terminus and non-amyloid beta component (NAC) region and an intrinsically disordered C-terminus that are predicted to increase its proclivity to phase separate (Mukherjee *et al*, 2023). *In vitro* experiments have shown that αSyn phase separates when there is increased molecular crowding (Ray *et al*, 2020), though the concentrations required are non-physiologically high. Post-translational modifications, mutations, pH changes, metal ions and binding partners such as the R-SNARE protein VAMP2 (synaprobrevin-2) and synapsin facilitate its phase separation (Agarwal *et al*, 2024; Dada *et al*, 2023; Hoffmann *et al*, 2021; Huang *et al*, 2022; Sawner *et al*, 2021; Wang *et al*, 2024; Xu *et al*, 2022). Aberrant LLPS of αSyn leads to a metastable porous hydrogel stage that eventually transitions into solid β-sheet-rich fibrillar aggregates (Hardenberg *et al*, 2021; Piroska *et al*, 2023; Ray *et al*., 2020). It has also been found that intermixed steps of LLPS and aggregation lead to the formation of αSyn inclusions (Eubanks *et al*., 2025).

Lipids are also involved in this process, in keeping with the nature of αSyn as a membrane-binding protein and with the fact that at least a proportion of Lewy bodies are rich in vesicles, membranes and lipids (Moors *et al*, 2021; Shahmoradian *et al*, 2019). Lipids influence the kinetics underlying the liquid-to-solid transition of αSyn: Liposomes, short saturated lipids and water soluble lipids arrest the maturation of αSyn condensates at the porous hydrogel stage, whereas long-chain lipids, cholesterol and fluid anionic membranes have the opposite effect (Galvagnion *et al*, 2016; Hardenberg *et al*., 2021).

Lipid droplets are connected to the dysregulation of αSyn. αSyn overexpression is linked to lipid droplet accumulation, and αSyn binding to lipid droplets leads to the formation of proteolysis-resistant αSyn aggregates (Girard *et al*, 2021; Smith *et al*, 2023). αSyn can also bind to the surface of lipid droplets and reduce the turnover of triglycerides (Cole *et al*, 2002). This connection could be particularly relevant to the pathogenesis of PD because midbrain dopaminergic neurons in the substantia nigra, which is one of the most vulnerable brain regions in PD, contain neuromelanin that is rich in lipid droplets (Filimontseva *et al*, 2025). Indeed, it has been shown that αSyn aggregates onto lipids present within neuromelanin and that changes in the lipids contained within neuromelanin precede the formation of Lewy bodies (Halliday *et al*, 2005).

Given these observations that aberrant LLPS, lipid dysregulation, lipid droplets and neuromelanin are all involved in αSyn dysregulation in PD, we propose an intriguing hypothesis: aberrant LLPS of αSyn upon binding to lipid droplets leads to the formation of hydrogel-like (semi-liquid/semi-solid) biomolecular condensates that are neurotoxic and cause neurodegeneration. In this manuscript, we tested this hypothesis, and our findings suggest a conceptual advance in our understanding of the pathogenesis of PD. We showed that αSyn condensates are cytotoxic and do not simply represent a metastable phase in the liquid-to-solid transition process. This toxicity occurs because they sequester lipid droplets and impede their turnover and consequent energy provision in the cell, as well as directly impair the function of neighbouring mitochondria.

## Results

### Lipid droplets facilitate the liquid-liquid phase separation of αSyn and the formation of biomolecular condensates

Recapitulating αSyn inclusions in cultured cells is a conundrum as none of the currently available model systems form Lewy body-like inclusions in a physiologically relevant manner. Seeding with recombinant αSyn pre-formed fibrils (PFFs) or human postmortem brain-derived fibrils leads to the formation of inclusions with no or limited ultrastructural similarities to Lewy bodies (Mahul-Mellier *et al*, 2020); in addition, the structure of recombinant PFFs does not accurately recapitulate that of αSyn fibrils from human brain (Todd *et al*, 2024). αSyn that carries three glutamic acid to lysine mutations in the KTKEGV repeat motifs in αSyn (E35K, E46K, E61K, “3K” αSyn) forms inclusions that ultrastructurally recapitulate a proportion of Lewy bodies (Moors *et al*., 2021; Shahmoradian *et al*., 2019), in that they are rich in vesicles, lipids and membranes that are surrounded by amorphously aggregated αSyn (Dettmer *et al*, 2015a; Dettmer *et al*, 2015b; Dettmer *et al*, 2017; Ericsson *et al*, 2021; Eubanks *et al*., 2025). However, the reliance on two artificial mutations (E35K, E61K) to “amplify” the effect of the Mendelian E46K mutation raises concerns about the true physiological relevance of this model.

We sought to develop a model system for αSyn inclusions that is more physiologically relevant and builds upon αSyn’s well characterized ability to aberrantly phase separate (Agarwal *et al*., 2024; Dada *et al*., 2023; Hardenberg *et al*., 2021; Hoffmann *et al*., 2021; Huang *et al*., 2022; Piroska *et al*., 2023; Ray *et al*., 2020; Sawner *et al*., 2021; Wang *et al*., 2024; Xu *et al*., 2022). Previous evidence suggested a potential relevance of lipid droplets to this process: It was recently reported that binding to lipid droplets promotes the phase separation of intrinsically disordered proteins (Kamatar *et al*, 2024). In addition, in our recent study we showed that 3K αSyn forms lipid droplet-rich inclusions, and that this ability is enhanced by the presence of increased numbers of lipid droplets (Eubanks *et al*., 2025). Finally, lipid droplets are enriched within the neuromelanin-rich dopaminergic neurons in the human substantia nigra, the most vulnerable brain region in PD (Filimontseva *et al*., 2025).

We treated M17D neuroblastoma cells stably overexpressing wild type (WT) or E46K mutant (1K) αSyn-YFP under doxycycline induction with Oleic Acid to induce the formation of lipid droplets (Papadopoulos *et al*, 2015). We selected the M17D cell line because it is neuronal and has previously been used to study αSyn dysregulation (Dettmer *et al*., 2017; Ericsson *et al*., 2021; Imberdis *et al*, 2019). Oleic acid treatment induced the formation of lipid droplet-rich αSyn inclusions at a ∼10-fold lower frequency as compared to 3K αSyn-YFP (**fig.1A,B**). The number of inclusions formed by E46K mutant or WT αSyn was proportional to the amount of time oleic acid was present in the culture media; the effect also diminished when oleic acid-containing media was replaced for 24h (**fig.1C,D,E**).

**Figure 1:**
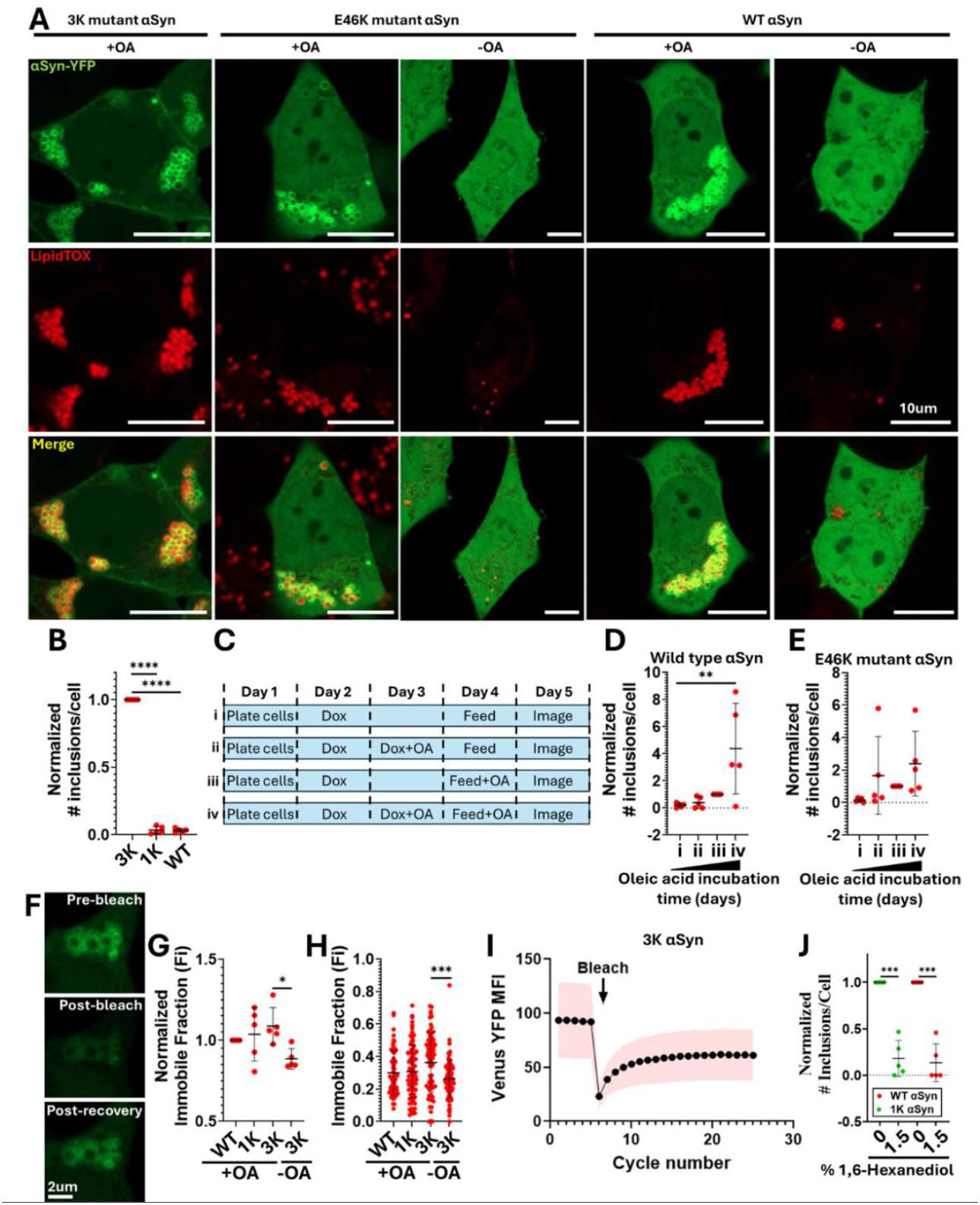
Lipid droplets induce the formation of αSyn inclusions. **A**. Confocal images of M17D cells expressing wild type (WT) or E46K mutant (1K) αSyn-Venus YFP after treatment with 600uM of oleic acid (OA) for 16h show the spontaneous formation of inclusions. Cells without oleic acid treatment did not form αSyn inclusions. Scale bar = 10um. **B**. 3K αSyn forms inclusions ∼10x more frequently than 1K and WT αSyn, all of which after treatment with 600uM of oleic acid. One sample t-tests. **C,D,E**. Directional increase in the number of αSyn condensates depending on the amount of time oleic acid was in the culture media. (C) describes each of the 4 conditions shown in (D) and (E). From left to right: i) No oleic acid; ii) oleic acid was added 1 day after doxycycline (Dox) induction, removed the following day and the cells were imaged the day after; iii) oleic acid was added 2 days after doxycycline induction and the cells were imaged the day after; iv) oleic acid was added 1 day after doxycycline induction, re-added the following day, and the cells were imaged the day after. One way ANOVA with test for linear trend. The results were significant for wild type (WT) αSyn (D) and there was a trend in the same direction for 1K αSyn (E). **F**. Representative timecourse FRAP images of a 1K αSyn inclusion. **G,H**. FRAP on WT, 1K and 3K αSyn inclusions with and without OA treatment. (G) shows the normalized data, whereas (H) is the unnormalized version of the same data. The no OA treatment condition applies only to 3K αSyn because it is the only one that spontaneously forms inclusions without lipid droplets. All conditions showed similar immobile fractions (Fi) indicating that they have similar liquid and solid proportions (liquid>solid). One way ANOVA with Tukey’s post-hoc correction. **I**. Representative FRAP recovery plot for 3K αSyn inclusions without OA treatment. The standard deviation (SD) is shown in pink. **J**. Treatment with 1.5% of 1,6-Hexanediol results in the significant reduction in the number of 1K and WT αSyn inclusions. One sample t-tests. All experiments were repeated independently 5 times, and the normalized values are shown in each plot, apart from (H) that contains the unnormalized pooled values from all independent experiments. Data are represented as mean +/-SD. P values: *≤0.05, **≤0.01, ***≤0.001, ****≤0.0001.

To ascertain the material status of the αSyn inclusions we undertook fluorescence recovery after photobleaching (FRAP) experiments which showed that WT and 1K αSyn inclusions had immobile fractions (Fi) of ∼0.35, similar to 3K αSyn (all of which after treatment with oleic acid). This indicates that these inclusions are largely liquid. Interestingly, 3K αSyn inclusions that formed spontaneously, without addition of oleic acid, were significantly more liquid than the more lipid droplet-rich 3K inclusions formed after treatment with oleic acid (**fig.1F,G,H,I**). Treatment with 1.5% 1,6-hexanediol significantly reduced the number of WT and 1K αSyn inclusions (**fig.1J**). These findings support that the lipid droplet-rich WT and 1K αSyn inclusions are consistent with biomolecular condensates.

### Lipid droplets modulate the number, type and S129 phosphorylation status of αSyn condensates

We sought to determine whether there is a dose response relationship between the oleic acid concentration in the culture media, the amount of lipid droplets and the number, size and type of αSyn condensates. We induced the formation of lipid droplets using two orthogonal methods: treatment with oleic acid or linoleic acid. We used the same concentrations and serial dilutions for each to enable robust comparisons of the results. Oleic acid and linoleic acid induce the formation of lipid droplets through separate mechanisms. Oleic acid is a mono-unsaturated omega-9 fatty acid; it is esterified and converted into triglycerides that incorporate into lipid droplets (Nakajima *et al*, 2019), or binds to the FFAR4 receptor on the cell surface that activates a signaling pathway including phosphoinositide 3-kinase and phospholipase D, which leads to the formation of new lipid droplets (Rohwedder *et al*, 2014). On the other hand, linoleic acid is a polyunsaturated, omega-6 fatty acid that is activated through esterification (Whelan & Fritsche, 2013) and incorporated into neutral lipids that are synthesized in the endoplasmic reticulum from which the lipid droplets eventually bud off.

We studied this process on 3K αSyn because it forms condensates much more frequently than 1K and WT αSyn, which facilitates their study. Higher concentrations of oleic acid and linoleic acid resulted in a significantly increased amount of lipid droplets as determined by the quantification of their surface area in relation to the total surface area of the cell within the same optical section; there was no difference between the same concentrations for these two treatments (**fig.2A**). This dose response effect was seen only for cells that contained αSyn condensates (**fig.2B**), which drove the results seen for the entire population of cells (**fig.2A**), but not for cells without (**fig.2C**). In addition, higher concentrations of linoleic acid and oleic acid correlated with increased fluorescence intensity of LipidTOX (staining neutral lipids within lipid droplets) (**fig.2D**). These results indicate that oleic acid and linoleic acid induce the formation of lipid droplets in a dose response manner and that there is a relationship between the amount of lipid droplets and the presence of condensates. In other words, it indicates that lipid droplets facilitate the formation of αSyn condensates.

**Figure 2:**
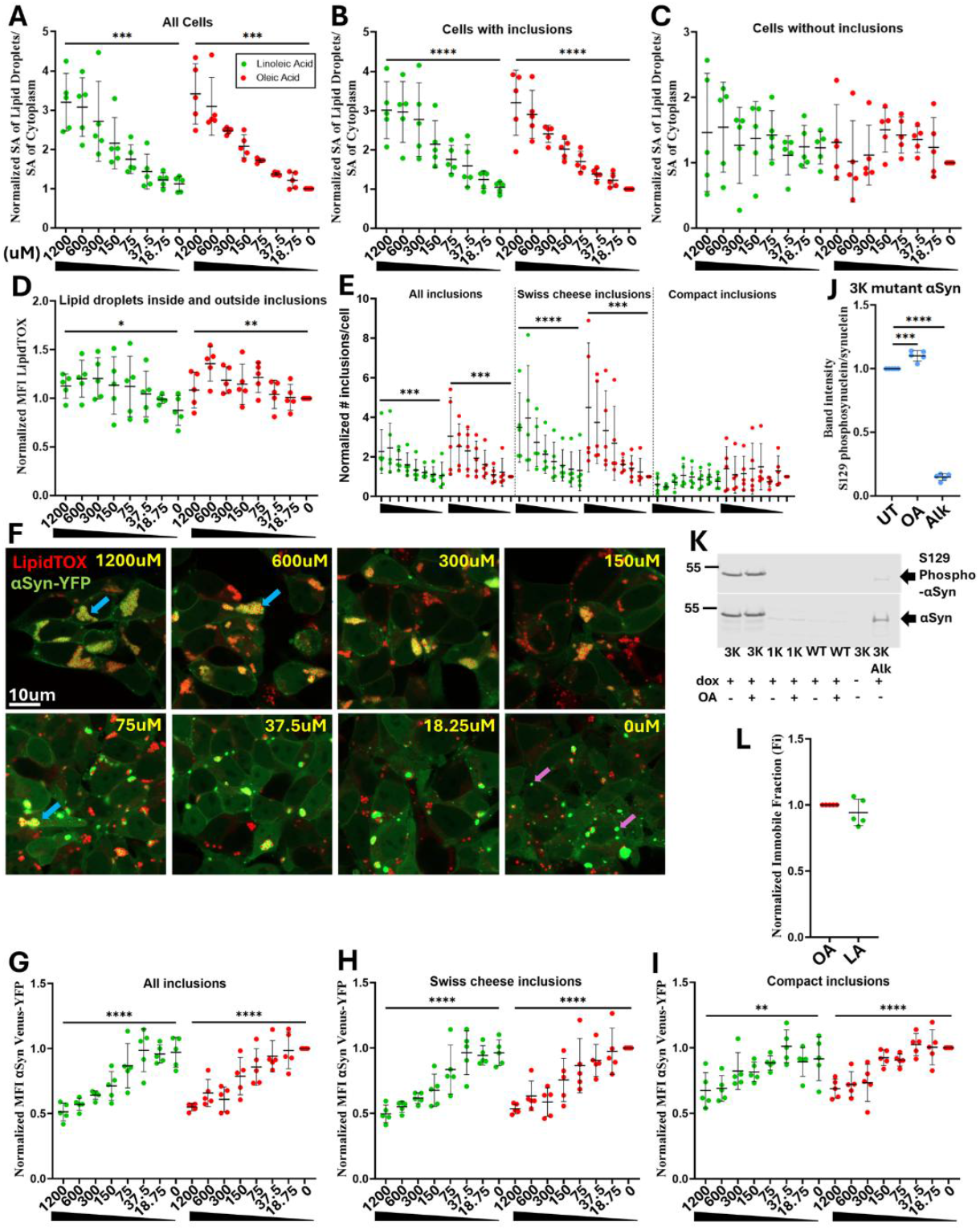
Lipid droplets modulate αSyn condensates. **A,B,C**. Dose response experiments for Linoleic acid (LA) and oleic acid (OA) showed that higher concentrations are significantly associated with an increased surface area (SA) of lipid droplets relatively to the total surface area of the cytoplasm, and that this association is driven by the cells containing αSyn condensates. The same concentrations were used in all dose response experiments for both lipids. One way ANOVA with test for linear trend. **D**. Dose response experiments for the mean fluorescence intensity (MFI) of the lipid droplets stained through LipidTOX far red showed that the MFI significantly increased as the concentrations of LA and OA increased. One way ANOVA with test for linear trend. **E**. Dose response experiments for LA and OA showed that the number of Swiss cheese condensates significantly increased with higher concentrations of LA and OA. There was no effect on the number of compact αSyn condensates. The same OA and LA concentrations were used as in (A,B,C). **F**. Representative confocal images for cells treated with the dose response of OA. Shown in blue arrows are Swiss cheese and in purple arrows compact αSyn condensates. **G, H, I**. The MFI of αSyn condensates (Swiss cheese, compact and all condensates) significantly decreased as the concentration of OA and LA increased. **J,K**. Western blot for S129 phosphorylated αSyn and total αSyn. 600uM of OA treatment for 24h at 2 days after doxycycline induction significantly increased the S129 phosphorylation of 3K αSyn. Alkaline phosphatase (Alk) was used as a positive control. 1K and WT αSyn show no phosphorylation consistent with our previous report (Eubanks *et al*, 2025). UT=cells without OA treatment. One way ANOVA with Tukey’s post-hoc correction. **L**. FRAP experiments for 3K αSyn αSyn formed after treatment with 600uM of LA or LA showed no difference in their immobile fractions (Fi). Concentrations in all plots are in uM. LA: green dots, OA: red dots. All experiments were repeated independently 5 times, and the normalized values are shown in each plot. Data are represented as mean +/-SD. P values: *≤0.05, **≤0.01, ***≤0.001, ****≤0.0001. SA, surface area.

To further corroborate the latter point, we quantified the number and type of αSyn condensates. As we previously showed, 3K αSyn condensates are of two different types: Swiss cheese-like that incorporate lipid droplets, and compact that do not (Eubanks *et al*., 2025). Our results indicate that both treatments significantly increase the total number of αSyn condensates and that this is driven by the increase in the number of Swiss cheese condensates; there is no difference seen for the compact condensates (**fig.2E,F**). Finally, higher concentrations of linoleic acid and oleic acid correlated with lower fluorescence intensity of αSyn-YFP within Swiss cheese and compact condensates (**fig.2G,H,I**). These results confirm that oleic and linoleic acid, and consequently lipid droplets, induce the formation of αSyn condensates and there is no difference in the efficiency associated with either treatment. The increase in the number of αSyn condensates is accompanied by decrease in their fluorescence intensity, indicating that the same total amount of αSyn is distributed among a larger number of condensates.

We assessed the S129 phosphorylation status of the αSyn condensates, as it is the most widely used marker for pathological αSyn and particularly αSyn aggregation. However, whether S129 phosphorylation induces aggregation, is a secondary event following aggregation, or inhibits further aggregation and toxicity is controversial (Ghanem *et al*, 2022; Oueslati, 2016). Treatment with oleic acid significantly increased the Fi of 3K αSyn condensates, thereby making them more solid (**fig.1G,H**). It also resulted in a small but significant increase in 3K αSyn S129 phosphorylation (**fig.2J,K, S1**). These findings suggest that increased lipid droplets induce the transformation of αSyn into a pathological S129 phosphorylated form. It is possible that lipid droplets act as a nidus on which αSyn binds (Eubanks *et al*., 2025) and congregates, resulting in its aggregation and secondary phosphorylation due to locally increased concentration.

FRAP experiments showed that the Fi of αSyn condensates formed after treatment with 600uM of either linoleic acid or oleic acid was the same (**fig.2L**). This indicates that comparable levels of lipid droplets, regardless of their path of induction and formation, result in the formation of αSyn condensates of similar material properties.

### αSyn condensates induce mitochondrial dysfunction

In our previous work we found that overexpression of 3K αSyn results in mitochondrial depolarization (Eubanks *et al*., 2025). It is well known that αSyn dysregulation affects mitochondrial function (Choi *et al*, 2022; Ludtmann *et al*, 2018). In this work, we observed that mitochondria surround αSyn condensates (**fig.3A**). Therefore, we hypothesized that αSyn condensates could impact mitochondrial function through direct contact. We used the 3K αSyn cells to test this hypothesis because the increased frequency of the condensates facilitated detection and robust analyses. After staining the live cells with tetramethylrhodamine methyl ester (TMRM) (in redistribution mode) and Hoechst, we undertook z-stack imaging and quantified the mitochondrial membrane potential (Δψ) as a general indicator of mitochondrial health in the vicinity of αSyn condensates (7-pixel radius) and in the remaining cytoplasm (Esteras *et al*, 2017). Mitochondria closely surrounding the αSyn condensates were significantly depolarized as compared to those further away in the cytoplasm (**fig.3B**). This suggests that αSyn condensates could directly induce mitochondrial dysfunction. Of note, this finding does not distinguish between lipid droplet-rich and -poor condensates; rather, it applies to αSyn condensates in general.

**Figure 3:**
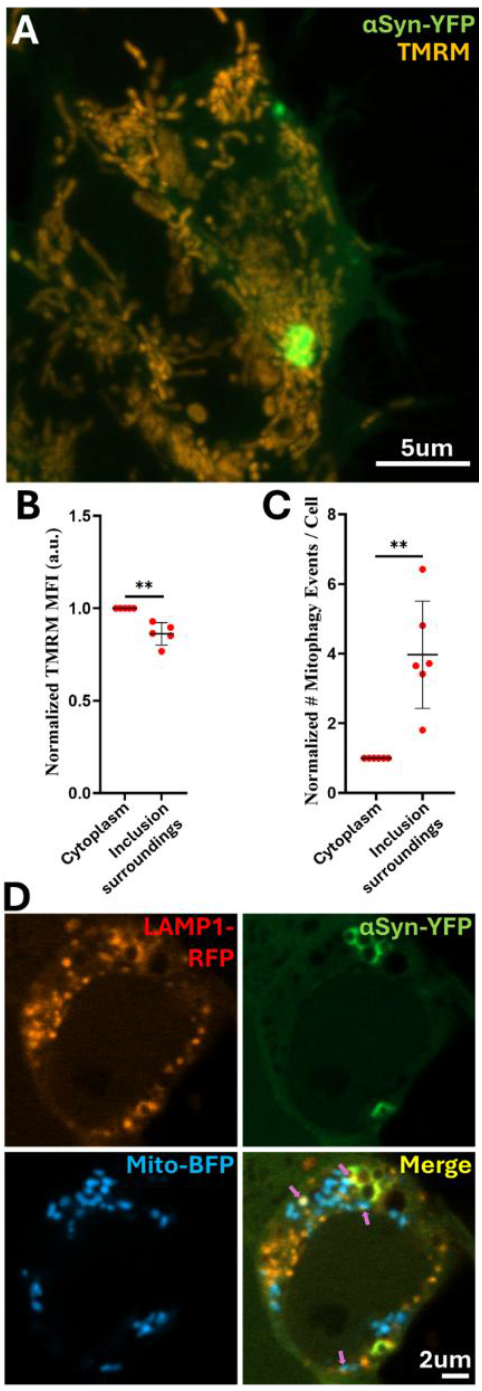
αSyn condensates impair mitochondrial function. **A**. Multiple intensity projection image generated from z-stack images of M17D cells expressing 3K αSyn-YFP showing that mitochondria closely approximate αSyn condensates. **B**. Mean fluorescence intensity (MFI) of tetramethylrhodamine methyl ester (TMRM) showing that mitochondria located within a 7-pixel radius around αSyn condensates are significantly depolarized. One sample t-test. a.u.=arbitrary units. **C,D**. Cells were transfected with a mito-BFP and LAMP1-RFP constructs to visualize the mitochondria and lysosomes, respectively. Late mitophagy events were determined by the colocalization of RFP with BFP. Mitophagy events were significantly more frequent within a 25-pixel radius around 3K αSyn inclusions than in the remaining cytoplasm. One sample t-test. All experiments were repeated independently 5 times, and the normalized values are shown in each plot. Data are represented as mean +/-SD. P values: *≤0.05, **≤0.01, ***≤0.001, ****≤0.0001.

We quantified mitophagy by determining the number of colocalization events between lysosomes and mitochondria, as indicated after transfection with LAMP1-RFP and mito-BFP constructs, respectively. This method identifies the number of mitophagy events independently of any specific pathways (Berezhnov *et al*, 2016). The number of mitophagy events was significantly increased around αSyn condensates as compared to the remaining cytoplasm (**fig.3C,D**). This indicates that mitochondria around αSyn condensates undergo increased removal.

### αSyn condensates entrap lipid droplets and impair their turnover

αSyn condensates frequently entrap lipid droplets as we showed earlier in this paper and in our previous work (Ericsson *et al*., 2021; Eubanks *et al*., 2025). An important function of lipid droplets is to store neutral lipids and release fatty acids through lipophagy for utilization by mitochondria when there are energy needs (Rambold *et al*, 2015; Singh *et al*, 2009). We hypothesized that the entrapment of lipid droplets in αSyn condensates impairs their turnover and energy provision in the cell.

We used a previously described pulse chase assay to quantify lipid droplet turnover (Rambold *et al*., 2015). In this assay, cells were pulsed with BODIPY 558/568 C12 overnight, which is a red fluorescent precursor to various phospholipids. On the following day they were chased overnight in regular culture media without dye, and on the day after they were stained with LipidTOX far red and imaged. This enables us to monitor the redistribution of C12 within the cell after it has been withdrawn from the culture media. We observed that the lipid droplets inside αSyn condensates were almost always double positive for C12 and LipidTOX, whereas those in the cytoplasm often lost the C12 staining. The same results were observed for 3K, 1K and WT αSyn (**fig.4A,B,C,D**). This indicates that lipid droplets that are free in the cytoplasm undergo a turnover process, but the ones entrapped in αSyn condensates do not.

**Figure 4:**
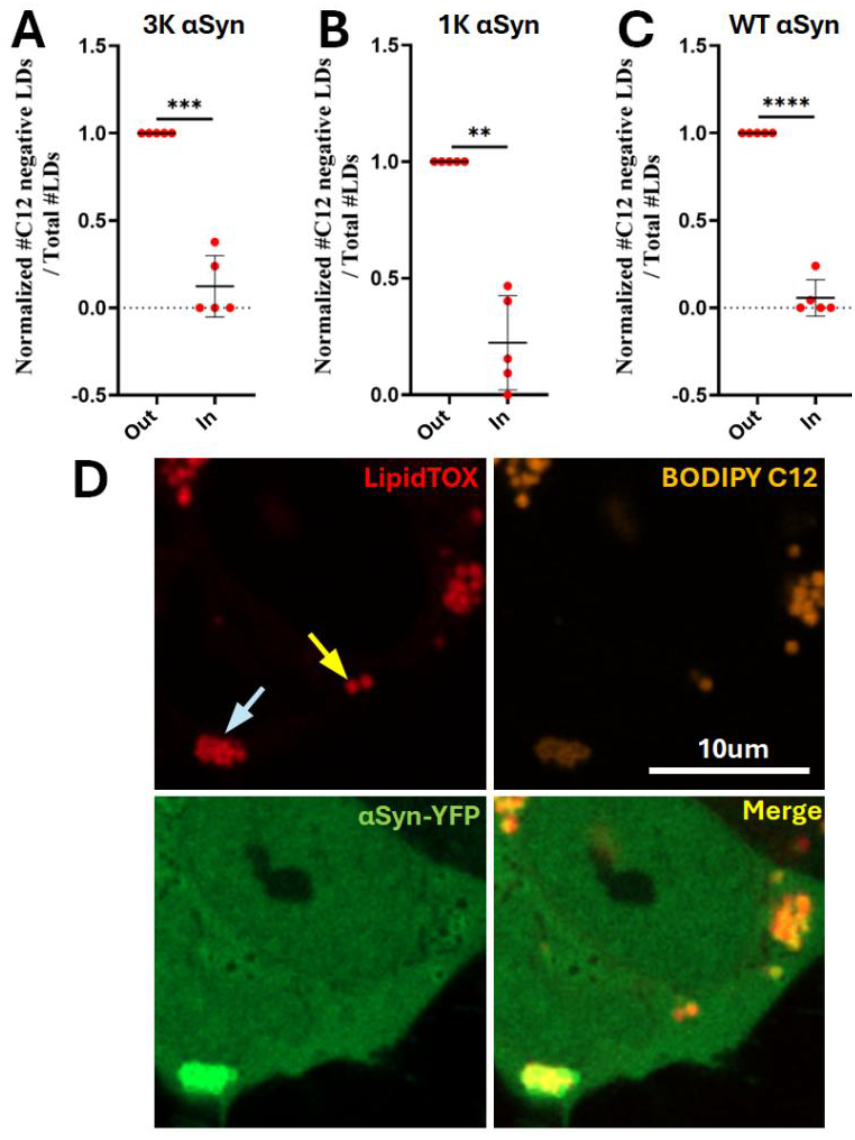
αSyn condensates entrap lipid droplets and reduce their turnover. **A,B,C**. Lipid droplets that are not entrapped in αSyn condensates (Out) show a loss of BODIPY 558/568 C12 staining more frequently than lipid droplets that are entrapped in 3K (A), 1K (B) and WT (C) αSyn condensates (In). Lipid droplets were identified through LipidTOX far red stain. One sample t-tests. **D**. Representative image of a WT αSyn condensate entrapping two LipidTOX-positive lipid droplets that are positive for BODIPY 558/568 C12 (blue arrow) and several cytoplasmic LipidTOX-positive lipid droplets, one of which is negative for BODIPY C12 (yellow arrow). All experiments were repeated independently 5 times, and the normalized values are shown in each plot. Data are represented as mean +/-SD. P values: *≤0.05, **≤0.01, ***≤0.001, ****≤0.0001.

## Discussion

There is robust but underexplored evidence suggesting that lipid droplets and aberrant LLPS are involved in the dysregulation of αSyn. This could be particularly relevant to the microenvironment of midbrain dopaminergic neurons in the substantia nigra given their contents in lipid droplet-rich neuromelanin. In a neuronal cell line, we found that αSyn aberrantly phase separates upon binding to lipid droplets and forms viscoelastic condensates (**fig.5A**) that impair energy provision in the cell by affecting lipid droplet turnover and mitochondrial function.

**Figure 5:**
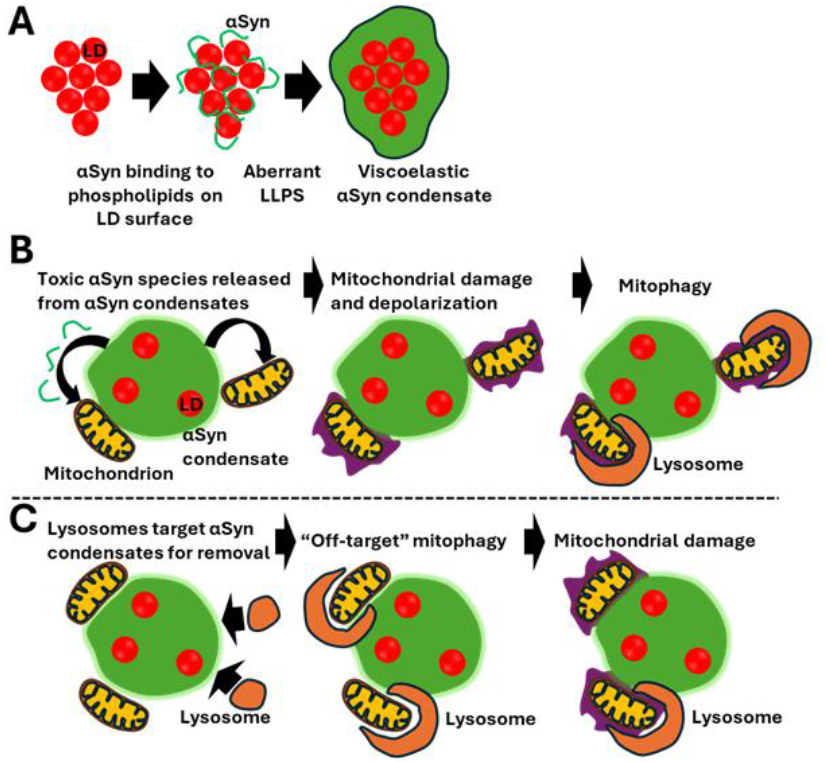
Proposed models for αSyn dysregulation in PD. **A**. Clusters of lipid droplets act as a nidus for αSyn binding, resulting in aberrant LLPS and the formation of viscoelastic condensates. **B,C**. Possible sequence of events connecting αSyn condensates to mitochondrial dysfunction. B. αSyn condensates release toxic species of αSyn due to their dynamic viscoelastic material status, which induce mitochondrial damage, as indicated by their alteration in membrane potential. The damaged mitochondria undergo mitophagy. C. Lysosomes target αSyn within condensates for removal, but also target nearby mitochondria as an off-target effect, which results in mitochondrial damage. LD, lipid droplets.

Our study indicates that biomolecular condensates are a toxic species of αSyn. αSyn condensates induce dysfunction of neighbouring mitochondria as indicated by their depolarization and increased mitophagy. It is possible that toxic species of αSyn bud off from the unstable, lipid-rich αSyn condensates and directly impact the integrity of the outer mitochondrial membrane. It is known that liposomes arrest the maturation of αSyn condensates at the hydrogel phase which acts as a reservoir for toxic oligomers (Hardenberg *et al*., 2021; Kumar *et al*, 2018). The sensitivity of mitochondria to toxic species of αSyn is well documented. αSyn converts from monomeric to oligomeric form upon interaction with mitochondrial membranes, attaches to cardiolipin and induces complex I dysfunction that leads to production of reactive oxygen species (Choi *et al*., 2022). αSyn oligomers also oxidize the ATP synthase beta unit, which induces lipid peroxidation and opening of the permeability transition pore and results in neurotoxicity (Ludtmann *et al*., 2018). The process itself leading to the formation of Lewy bodies is toxic for mitochondria (Mahul-Mellier *et al*., 2020). Interestingly, a similar mechanism has been reported for Huntington’s disease: polyglutamine (polyQ)-expanded huntingtin inclusions directly interact with mitochondrial membranes, which leads to mitochondrial dysfunction (Bauerlein *et al*, 2017). This indicates that mitochondrial dysfunction that is induced through direct interaction with proteinaceous inclusions could be a generalizable model for the pathogenesis of neurodegenerative diseases. Studies on postmortem brain tissue support the neurotoxicity of Lewy bodies: neurons containing Lewy bodies exhibit microtubule regression and mitochondrial degradation (Power *et al*, 2017).

αSyn condensates entrap lipid droplets impeding their turnover, as indicated by our BODIPY C12-LipidTOX pulse chase assay. It is known that lipid droplets store fatty acids as triacylglycerols which can be released for energy production purposes either through lipophagy that entails engulfment of the lipid droplet by autophagosomes and fusion with lysosomes, or through direct hydrolysis by lipases (lipolysis). Mitochondria take up the fatty acids that are released through lipolysis where they undergo β-oxidation, a process that requires lipid droplets and mitochondria to be in close proximity (Rambold *et al*., 2015). Therefore, the sequestration of lipid droplets in αSyn condensates could prevent the provision of critical fatty acids to mitochondria, leading to their dysfunction.

Mitochondria surrounding αSyn condensates undergo increased mitophagy. It is possible that this occurs as a secondary phenomenon to their dysfunction that is caused by either the direct impact of αSyn condensates or inadequate fatty acid provision from lipid droplets (**fig.5B**). Alternatively, it is possible that it is an “off target” effect: lysosomes could be trying to access and digest αSyn condensates but instead target the mitochondria that just happen to be in the way (**fig.5C**).

Our study describes a new, physiologically relevant model system for αSyn inclusions that is based on aberrant LLPS: inducing the formation of lipid droplets through treatment with oleic acid leads to the spontaneous formation of condensates by WT and E46K mutant αSyn. It is likely that this model system recapitulates the early stages of αSyn inclusions, given several orthogonal reports that implicate the early involvement of aberrant LLPS in their formation process (Agarwal *et al*., 2024; Hardenberg *et al*., 2021; Piroska *et al*., 2023; Ray *et al*., 2020; Wang *et al*., 2024). Our findings also support the physiological relevance of the 3K αSyn model system which has been questioned because it involves two artificial mutations (E35K and E61K). Nevertheless, 3K αSyn does form inclusions with striking ultrastructural similarities to Lewy bodies (Eubanks *et al*., 2025; Moors *et al*., 2021; Shahmoradian *et al*., 2019). In this study, we also found that the 3K αSyn condensates are similar in their behaviour and functional consequences to WT and E46K mutant αSyn condensates, with the only difference being that 3K αSyn forms ∼10x more frequent condensates, presumably due to its enhanced interactions with phospholipids on lipid droplets due to the extra positively charged lysines (Eubanks *et al*., 2025).

Our findings underscore the relevance of lipid droplets to neurodegeneration and αSyn dysregulation in PD. Lipid droplets have been implicated in the neurodegeneration process. Neurons under oxidative stress actively secrete peroxidated lipids that are taken up by microglia through fatty acid transporters and Apolipoprotein E and accumulate in the form of lipid droplets (Liu *et al*, 2017; Liu *et al*, 2015; Moulton *et al*, 2021). Lipid droplets have also been extensively implicated in the pathogenesis of hereditary spastic paraplegias, which is supported by the fact that many genes involved in lipid droplet homeostasis carry Mendelian mutations in these syndromes (Kara *et al*, 2016). Lipid droplets are particularly enriched in neuromelanin-containing midbrain dopaminergic neurons in the substantia nigra, which degenerate early on in PD. It is therefore entirely plausible that this unique microenvironment facilitates the αSyn-associated early degeneration of the substantia nigra in PD. Lewy bodies exhibit structural heterogeneity in terms of their protein and lipid composition (Lashuel, 2020; Shahmoradian *et al*., 2019; Wakabayashi *et al*, 2007). It is possible that lipid-rich αSyn condensates are uniquely enriched in the substantia nigra, which develop into lipid-rich Lewy bodies with arrested maturation (Hardenberg *et al*., 2021) that maintain continued cytotoxic properties (Kumar *et al*., 2018) leading to early neurodegeneration.

## Methods

### List of reagents

| REAGENT OR RESOURCE | SOURCE | IDENTIFIER |
| --- | --- | --- |
| **Antibodies** |  |  |
| Rabbit anti-α-synuclein (phosphoS129) | Abcam | Cat #:ab168381 |
| Mouse anti-α-synuclein | BD Biosciences | Cat #:610786 |
| IRDye 680RD goat anti-rabbit | LiCor | Cat #:925-68071 |
| IRDye 800CW goat anti-mouse | LiCor | Cat #:925-32210 |
| **Chemicals, peptides, and recombinant proteins** |  |  |
| Dulbecco's Modified Eagle Medium (DMEM) | Thermofisher | Cat #:11965118 |
| Fetal Bovine Serum (FBS) | Thermofisher | Cat #:16000044 |
| Glutamax | Thermofisher | Cat #:35050061 |
| Penicillin-Streptomycin | Thermofisher | Cat #:15140122 |
| Trypsin | Thermofisher | Cat #:25300054 |
| Dulbecco's Modified Eagle Medium (DMEM) without phenol | Thermofisher | Cat #:31053028 |
| Oleic Acid-Albumin from bovine serum | Sigma | Cat #:O3008-5mL |
| 8 well coverglass bottom chamberslides | Ibidi | Cat #:80807 |
| doxycycline hyclate | Sigma | Cat #:D9891-1G |
| Opti-MEM reduced serum medium (OMEM) without phenol | Thermofisher | Cat #:11058021 |
| Lipofectamine RNAiMAX Transfection Reagent | Thermofisher | Cat #:13778150 |
| Lipofectamine™ 3000 Transfection Reagent | Thermofisher | Cat #:L3000015 |
| Hoechst 33342 trihydrochloride trihydrate nuclear dye | Thermofisher | Cat #:H3570 |
| 1,6-Hexanediol | Sigma | Cat #:240117-50G |
| Sterile distilled water | Gibco | Cat #:15230162 |
| HCS LipidTOX™ Deep Red Neutral Lipid Stain | Thermofisher | Cat #:H34477 |
| Tetramethylrhodamine, Methyl Ester, Perchlorate (TMRM) | Thermofisher | Cat #:T668 |
| Dulbecco's phosphate-buffered saline (DPBS), calcium, magnesium | Gibco | Cat #:14040133 |
| Dulbecco's phosphate-buffered saline (DPBS), no calcium, no magnesium | Thermofisher | Cat #: 14190144 |
| Phosphate buffered saline (PBS) | Sigma | Cat #:P3813-10PAK |
| Linoleic Acid-Albumin from bovine serum albumin | Sigma | Cat #: L9530-5ML |
| RIPA Buffer (10X) #9806 | Cell Signaling | Cat #: 9806S |
| Protease inhibitor cocktail | Sigma | Cat #: 11697498001 |
| Halt™ Phosphatase Inhibitor Single-Use Cocktail | Thermofisher | Cat #: 78420 |
| FastAP Thermosensitive Alkaline Phosphatase (1 U/μL) | Thermofisher | Cat #: EF0651 |
| Pierce BCA Protein Assay Kit, 500 mL | Thermofisher | Cat #: 23227 |
| NuPAGE LDS Sample Buffer, 4X (NP0007) | Thermofisher | Cat #: NP0007 |
| NuPAGE Sample Reducing Agent, 10X (NP0009) | Thermofisher | Cat #: NP0009 |
| NuPAGE™ 4 to 12%, Bis-Tris, 1.0–1.5 mm, Mini Protein Gels | Thermofisher | Cat #: NP0321BOX |
| Amersham Protran Premium 0.2 NC 150mm×4m 1 roll/PK | Cytiva | Cat #: 10600014 |
| Odyssey Blocking Buffer | LiCor | Cat #: 927-60001 |
| NuPAGE MES SDS Running Buffer, 20X (NP0002) | Thermofisher | Cat #: NP0002 |
| PageRuler Plus Prestained Protein Ladder, 10 to 250 kDa (26619) | Thermofisher | Cat #: 26619 |
| Intercept Antibody Diluent | LiCor | Cat #: 927-65001 |
| Fisher BioReagents™ Microbiology Media: LB Agar, Miller (Powder) | Fisher Scientific | Cat #:BP1425-500 |
| Fisher BioReagents™ Microbiology Media: LB Broth, Miller | Fisher Scientific | Cat #:BP1426-500 |
| Ampicillin | Sigma | Cat #:A0166-5G |
| Kanamycin | Sigma | Cat #:K1377-1G |
| FastAP Thermosensitive Alkaline Phosphatase (1 U/μL) | Thermofisher | EF0651 |
| BODIPY 558/568 C12 | Thermofisher | Cat #: D3835 |
| BODIPY 493/503 | Thermofisher | Cat #: D3922 |
| One Shot™ MAX Efficiency™ DH5α-T1R Competent Cells | Thermofisher | Cat #:12297016 |
| **Critical commercial assays** |  |  |
| HiSpeed Plasmid Maxi Kit (25) | Qiagen | Cat #:12663 |
| MycoAlert® PLUS Mycoplasma Detection Kit (30 Tests) | Lonza | Cat #:LT07-703/<br>NC0529908 |
| **Experimental models: Cell lines** |  |  |
| M17D 3K αSyn cell line, male origin | Drs. Ulf Dettmer and Tim Bartels (Dettmer *et al.*, 2017) | N/A |
| M17D E46K (1K) αSyn cell line, male origin | Drs. Ulf Dettmer and Tim Bartels | N/A |
| M17D wild type αSyn cell line, male origin | Drs. Ulf Dettmer and Tim Bartels | N/A |
| **Recombinant DNA** |  |  |
| Mito-BFP construct | Addgene | Cat #: 49151 |
| Lamp1-RFP construct | Addgene | Cat #: 1817 |

**Note:** The above list of reagents significantly overlaps the list recently published by us (Eubanks *et al.*, 2025) due to the methodological overlap of the studies.

### Cell line generation

Synthetic human aS-WT::YFP and aS-E46K::YFP cDNA fragments were digested with BamHI and EcoRI and inserted into the corresponding restriction sites of the pLVX-TetOne-Puro lentiviral vector (Clontech/TaKaRa, Mountain View, CA). Silent mutations were introduced to BamHI and EcoRI recognition sites within the *SNCA* coding sequence. The recombinant plasmid was sequence-verified and subsequently used for lentiviral particle production, following previously described protocols (Dettmer *et al*., 2017). Early-passage human M17D neuroblastoma cells were transduced with aS-WT::YFP or aS-E46K::YFP lentiviral particles and selected for stable integration using puromycin. Cells expressing the YFP-tagged transgene were further enriched by fluorescence-activated cell sorting (FACS) after doxycycline induction. Resulting cell pools were expanded, tested again for expression upon dox induction, and absence of mycoplasma was confirmed.

### Tissue culture

#### Cell maintenance

M17D neuroblastoma cells stably overexpressing E35K + E46K + E61K mutant (3K) αSyn, E46K mutant (1K) or wild type (WT) αSyn fused with Venus YFP on their C-terminus, kindly shared by Profs Ulf Dettmer and Tim Bartels were used in this study. Cells were maintained in Dulbecco’s Modified Eagle Medium (DMEM) (Thermofisher, Cat #:11965118) supplemented with 10% fetal bovine serum (FBS) (Thermofisher, Cat #: 16000044), 1% Penicillin-streptomycin (Thermofisher, Cat #:15140122) and 1% Glutamax (Thermofisher Cat #:35050061). Cells were kept at 37°C, >90% humidity and 5% CO_2_. At 70-80% confluency, the cells were trypsinized (Thermofisher Cat #:25300054) and seeded in new flasks. Cells were periodically tested for mycoplasma contamination with the MycoAlert PLUS Mycoplasma Detection Kit (Lonza, Cat #:LT07-703). For imaging experiments, the cells were switched to DMEM without phenol red (Cat #:31053028) supplemented with FBS, glutamax and penicillin-streptomycin as described above, after plating.

#### Cell plating

Cells were seeded in 8 well chamber slides or 6 well plates depending on experimental needs, at densities of 2.2x10^5^ cells/mL. 300uL/well and 2mL/well were seeded in the 8 well chamberslides and 6 well plates, respectively.

#### Doxycycline induction

24h after seeding, cells were treated with 1ug/mL doxycycline hyclate (#D9891-1G, Sigma) in DMEM without phenol supplemented with FBS, penicillin/streptomycin and glutamax, to induce expression of αSyn-YFP. 48h later, the media was switched to media without doxycycline. Experiments were conducted either on the same day or later.

#### 1,6-Hexanediol

Stock 1,6-Hexanediol solution was prepared as we previously described (Eubanks *et al*., 2025). 1.5% concentration that was incubated for 30min was used in our experiments.

### αSyn inclusion quantification through tiling images

For each condition we acquired 3x3 tiling images at 20x for the YFP channel with 488nm excitation. We segmented whole cells using Cellpose. Within each labeled cell, we detected inclusions using adaptive Otsu thresholding. For each cell we recorded the count of inclusions. Results were exported to Excel tables.

### αSyn inclusion quantification through 63x images

3K αSyn cells were treated with doxycycline. 48h later they were treated with dose-response concentrations of Oleic acid-albumin (Sigma Cat #:O3008-5mL) and Linoleic acid-albumin (Sigma Cat #: L9530-5ML). The concentration of the stock solution was determined through the specific lot number of the bottle as per manufacturer instructions. An initial concentration of 1200uM was prepared in DMEM without phenol red+FBS+penstrep+glutamax, which was then used for serial dilutions: 600, 300, 150, 75, 37.5, 18.75uM. An untreated control (0uM) was also used. Lipid droplets were visualized by staining with LipidTOX deep red (Thermofisher Cat #:H34477) at a concentration of 1:500 for 1h.

We quantified the number and types (Swiss cheese, compact) of αSyn inclusions, MFI of αSyn inclusions, MFI and surface area of lipid droplets and the surface area of the cytoplasm through Python code. Micro SAM is a finetuned version of SAM (Segment Anything Model). Micro SAM extends SAM by finetuning for microscopy images to segment cells, organelles, and other objects typically found in biological images. We finetuned the pretrained vit-b-lm model on 442 annotated images to segment the αSyn inclusions to improve segmentation accuracy. We used 48 images for testing to evaluate the improvement in accuracy from the previous process which resulted in an improvement from 40% to 70% using the IoU metric. The code separates the image into its two separate channels, YFP and deep red. It starts by first preprocessing the YFP channel by applying a Gaussian blur to reduce noise, enhancing contrast using a sigmoid adjustment, and normalizing the image from 0 to 1. This image is then input into the pretrained machine learning model to segment the inclusions. Next, the code generates a labelled image of the cells from the YFP channel using cellpose. The lipid droplets are segmented into their own masks using an intensity threshold. The code then does a loop through each cell in the labelled cell mask and extracts the inclusions and lipid droplets in each cell. It then classifies each inclusion as either Swiss cheese or compact by checking if it has any overlap with a lipid droplet. Afterwards, it counts how many inclusions are in each cell, how many Swiss cheese or compact, and the lipid droplets in either the cytoplasm or inclusions. To measure the Mean Fluorescence Intensity (MFI), the code performs segmentation of inclusions, cells, and lipid droplets from the green and deep red channels. It loops through each cell in the labeled cell mask and extracts the inclusions and lipid droplets that overlap with each cell. Inclusions are then classified as either Swiss cheese or compact, and lipid droplets are categorized based on whether they are located inside or outside the inclusions. Using these segmented masks, the code retrieves the corresponding regions from the original image to access pixel intensity values. The MFI is calculated by summing the fluorescence signal within a mask and dividing by the number of pixels in that mask. This process is repeated to compute the MFI of lipid droplets and αSyn inclusions. Finally, the computed data for each cell is compiled and exported into an Excel spreadsheet.

### Mitophagy experiments

#### Transfections

The following constructs were used for transfections: mito-BFP was a gift from Gia Voeltz (Addgene plasmid # 49151 ; http://n2t.net/addgene:49151 ; RRID:Addgene_49151) (Friedman *et al*, 2011). Lamp1-RFP was a gift from Walther Mothes (Addgene plasmid # 1817 ; http://n2t.net/addgene:1817 ; RRID:Addgene_1817) (Sherer *et al*, 2003). Cells were transfected 48h after doxycycline induction and imaged 2 days later. RNAiMAX (#13778150, Thermofisher) was used for the 3K M17D cells as we previously described (Eubanks *et al*., 2025). Lipofectamine 3000 (Cat #:L3000015) was used for the 1K and WT M17D cells because of reduced transfection efficiency with RNAiMax.

Mitophagy events were determined by the colocalization between mito-BFP and LAMP1-RFP puncta, as analyzed through Python code. First, it separates the image into its three separate channels (BFP, YFP, RFP). Then it preprocesses the green channel by applying a Gaussian blur to reduce noise, enhancing contrast using a sigmoid adjustment, and normalizing the image from 0 to 1. This image is then input into a pretrained machine learning model to segment the inclusions from the green images. Next, the code generates a labeled image of the cells from the YFP channel using cellpose. The orange and blue channels’ desired objects are then segmented using an intensity threshold. The orange thresholding mask is then filtered to exclude signal overlapping with the YFP channel, which represents either bleedthrough or true signal due to incorporation of lysosomal fragments in αSyn inclusions (Dettmer *et al*., 2017). The YFP mask is then dilated by a radius of 25 pixels. This mask is then cut down by removing the original inclusion objects, resulting in a “donut” mask surrounding the inclusions. The code then loops through each cell within the labelled cell mask. It extracts the YFP+ inclusions, RFP+ lysosomes, and BFP+ mitochondria within each cell, creating a separate mask for each of those. It also obtains the “donut” mask within each cell and extracts the BFP+ and RFP+ objects specifically within the donut mask region and outside the dilated inclusion region. Finally, it counts the mitophagy events in the donut region and outside the dilated area by counting how many RFP+ objects touch or overlap with a BFP+ object. This data is then exported as an Excel datasheet.

### Pulse chase assay for lipid droplet turnover

This assay was undertaken as previously described (Rambold *et al*., 2015). Cells were stained overnight with 1µM of BODIPY 558/568 C12 (Thermofisher, Cat #: D3835), which is a fluorescent precursor to various phospholipids. They were then chased in DMEM without phenol red + FBS + glutamax + penstrep for 24h, which was supplemented with 600uM of oleic acid for the 1K and WT cells. This was followed by staining with LipidTOX deep red (Thermofisher Cat #: H34477) 1:500 for 1h and then by imaging. BODIPY 558/568 C12 was excited with the 561nm laser, YFP with 488nm and LipidTOX deep red with 640nm.

Data was analysed through Python code. First, it separates the image into its three separate channels (LipidTOX deep red, C12, and YFP). It starts off by first preprocessing the YFP channel by applying a Gaussian blur to reduce noise, enhancing contrast using a sigmoid adjustment, and normalizing the image from 0 to 1. This image is then input into a pretrained machine learning model to segment the inclusions from the YFP images. Next, the code generates a labelled image of the cells from the YFP channel using Cellpose. The deep red and C12 objects are segmented into their own masks using an intensity threshold. Then a separate mask is created for each of the deep red and C12 channels. If an object in the deep red mask is not overlapping or touching with a C12 mask, then it is added to the deep red-only mask, which represents the lipid droplets that do not exhibit C12 staining. The code then loops through each cell within the labelled cell mask. Within each cell, it extracts the inclusions. Then, for each individual cell, the code analyses the spatial relationship between inclusions and lipid droplets across different masks. For each category (total, deep red-only), the script determines how many lipid droplets are located within inclusions versus outside.

### Fluorescence recovery after photobleaching (FRAP)

FRAP experiments were undertaken using either a Zeiss LSM880 confocal with AiryScan with a 63x oil-immersion objective or a Zeiss LSM900 confocal with AiryScan with a 63x oil-immersion objective as we previously described (Eubanks *et al*., 2025). Cells were imaged 48h after doxycycline induction. For the comparison between linoleic acid and oleic acid-treated 3K cells, treatment with 600uM of each lipid for 16h was used. Laser settings (power and number of frames) were optimized to ensure less than 10% background bleaching during the timecourse experiment. A region of interest (ROI) was drawn around the αSyn inclusion of interest. For 3K αSyn inclusions, 5 images were taken pre-bleach, followed by bleaching at 100% transmission with the 488nm laser for 8 iterations and 35 post-bleaching cycles. For 1K and WT αSyn inclusions, 3 images were taken pre-bleach, followed by 12 images post-bleach. Of note, the total duration of the timecourse was the same in all these experiments. For analysis, ROIs were drawn for the cytoplasm within the same cell and the black background outside cells. All downstream steps, described in detail in our previous public publication (Eubanks *et al*., 2025) were undertaken through a Python script.

First, the data from the three ROIs are imported into the script through an Excel sheet. On top of this, the script requires some preliminary data from the user, such as the time of the bleaching and the timeframe. Every number in the columns is truncated to two decimal places. With the truncated data, it now computes several data points, such as the average numbers in each ROI before being bleached, and specifically for ROI2 (cytoplasm), it calculates the average of the numbers after being bleached. With these values, the script proceeds to calculate the normalized fluorescence intensity for ROI1. This is done by subtracting the corresponding ROI3 value (background) from the ROI1 value and dividing the result by the post-bleach average of ROI2. This normalized value represents the fluorescence recovery trend over time and is recorded in a new column. The script then fits these normalized values to an exponential recovery model. Once it fits, the code extracts an important parameter from the model, which is the plateau value. Using the plateau and the average of the normalized values before bleaching, it further calculates the mobile fraction and the immobile fraction. All of the data is exported as an Excel spreadsheet. After this, a smooth exponential recovery curve is generated based on the fitted parameters from the FRAP analysis. It creates 1000 evenly spaced time points from 0 to the final recorded time, and uses the fitted exponential model to compute the corresponding fluorescence intensity values for each time point. These values are stored in a new DataFrame along with their normalized percentage values. The DataFrame is then exported as an Excel. Finally, the script generates a graph for the raw data, plotting the numbers of all three ROIs over time. A second graph is produced for the normalized values of ROI1, showing both the actual data points and the fitted exponential recovery curve. Lastly, a third graph is created to display the fitted curve in terms of percentage recovery, with a dashed line marking 100% as a reference for complete fluorescence restoration. All three plots are formatted for clarity and saved together as a single .jpg image to provide a comprehensive visual summary of the FRAP analysis. Quality control for the analysis of each individual inclusion was undertaken manually.

For the experiment where the liquid status of αSyn inclusions near and far from mitochondria was studied, the former was determined through visual inspection depending on whether or not the inclusion on which we conducted FRAP was touching any mitochondria.

### Mitochondrial membrane potential measurement

Cells were incubated for 40min with 1μg/mL Hoechst 33342 (Thermofisher Cat #:H3570) and 100nM tetramethylrhodamine methyl ester (TMRM) (Thermofisher Cat #:T668) at 48h after doxycycline induction as we previously described (Eubanks *et al*., 2025). Z-stacks were acquired for the Hoechst (excitation 405nm), YFP (excitation 488nm) and TMRM (excitation 561nm) channels.

For analysis, cells were segmented through Cellpose (cyto3 model) as follows. The YFP and nuclear channels were stacked, we ran the 2D model on each slice and then enabled cross-stitching to link masks with sufficient overlap between adjacent slices. To accommodate size variability, we swept the diameter parameter and returned the first non-empty segmentation. Nuclei were segmented through Cellpose. A mitochondria binary mask was derived from the orange channel using Otsu thresholding. Inclusions were detected independently within each labeled cell and slice from the green channel using an adaptive threshold method. Then we filtered the inclusions to keep the ones with area 10-10,000 pixels, then we removed the ones with circularity index <= 0.1. For each cell we defined 4 ROIs: Cell, Inclusions, Inclusion surroundings (dilation radius = 7), and Cell without Inclusions and Surroundings. For each ROI per slice, we calculated the MFI = mean raw TMRM intensity inside the thresholded mitochondria binary mask in the ROI. Values were averaged across the z axis to yield per-cell results.

### Western blot

3K, 1K and WT cells were plated at 2.2x10^5^ cells/mL density in 6 well plates. After doxycycline induction for 48h, the cells were treated with 600uM of oleic acid for 16h. They were then trypsinized and pelleted. The pellet was lysated for 15min on ice through RIPA buffer (CellSignaling, Cat#: 9806S) containing protease (Sigma, Cat#: 11697498001) and phosphatase inhibitors (Thermofisher, Cat#: 78420). The latter was omitted in the sample that would be used for the Alkaline phosphatase treatment (positive control). The samples were centrifuged at 10,000g at 4°C for 15min. The protein concentration of the supernatant was measured through BCA assay (Thermofisher, Cat#: 23227). The samples were mixed with LDS (Thermofisher, Cat#: NP0007) and reducing reagent (Thermofisher, Cat#: NP0009), ran on 4-12% Bis-Tris gels (Thermofisher, Cat#: NP0321BOX), transferred onto nitrocellulose membranes (Cytiva, Cat#: 10600014) overnight, blocked with Odyssey blocking buffer (LiCor, Cat#: 927–60001) and incubated with the primary antibody diluted in Intercept antibody diluent (LiCor Cat#: 927–65001) overnight. The following day, the membranes were washed with TBST 4 times and TBS 1 time, for 10min each. Secondary antibodies were incubated for 1-2h at room temperature. The membranes were washed and imaged on a LiCor Odyssey CLx. The following antibodies were used: Mouse anti-αSyn 1:2000 (BD Biosciences Cat#: 610786), Rabbit anti-α-synuclein (phosphoS129) 1:1000 (Abcam, Cat#: ab168381). IRDye 680RD goat anti-rabbit 1:2000 (LiCor, Cat#: 925-68071), IRDye 800CW goat anti-mouse 1:2000 (LiCor, Cat#: 925-32210). Images were analyzed in ImageJ.

### Statistical analysis

Statistical analyses were undertaken using GraphPad Prism version 10.3.0. Mean +/-standard deviation (SD) is shown on all plots. Statistical tests used include one sample t-test, one way ANOVA with post hoc Tukey’s correction for multiple testing, and one way ANOVA with test for linear trend. Correction for multiple testing was undertaken as appropriate. After optimization, each experiment was repeated independently 5 times. The data was normalized within each experiment to a common control condition, followed by pooling the data and statistical analysis, unless otherwise stated in the figure legends. Only significant differences are indicated (with stars) on the graphs. The statistical test, number of biological replicates and significance levels are indicated in the respective figure legends. The images shown in the figures of this paper are after contrast adjustment that was applied uniformly to the entire channel. Analysis of the images through Python code was done as described in the methods section.

## Data accessibility

Python code used in data analysis in this manuscript has been posted publicly on GitHub: Kara-Lab/analyses_of_alpha-synuclein_biomolecular_condensates at main · eleannakara/Kara-Lab · GitHub.

## Acknowledgments

We acknowledge the following funding for student research fellowships: NIH R25 grant (R25NS105143) to Rutgers NeuroSURP summer student program that funded a student in our lab (AM), AOA Carolyn L. Kuckein Student Research Fellowship (EE), Jack Kent Cooke foundation graduate scholarship (JC), Aresty summer science research fellowship (NRS), Rutgers Health Center for Biomedical Informatics & Health Artificial Intelligence (BMIHAI) summer research internship (SJ). This work was funded through Rutgers start-up funding (EK). We are grateful to Katelyn VanderSleen, Neha Patel, Philip Socha and Sreenidhi Ravishankar for technical assistance during early stages of this project, and Heba Alnakhala for technical assistance with generating the M17D-based cell lines.

## Author contributions

Study conception and design: EK

Study supervision: EK

Funding acquisition: EK

Generation of cDNA constructs, viral particles and cell lines: NRam, AT, UD

Tissue culture: EG, JC, EE, NRS, AD, NR, AM

FRAP experiments and data analysis: JC, EE, EG, NRav, AD, AM, NRS

Mitophagy experiments: JC

Mitochondrial experiments: JC, EE

Pulse chase experiments: JC, EE, EG

Imaging of αSyn inclusions: EE, EG, JC, NRS, NRav, AD, AM

Performed image annotations: NRS, AD, AM, EG

Python code: SJ, YH

Data analysis: JC, EE, SJ, YH

Wrote paper: EK, EE, JC, SJ, YH

Edited paper and provided comments: EK, UD, EE

Provision of critical reagents and advice: UD, TB

## Supplementary figures

**Supplementary Figure 1:**
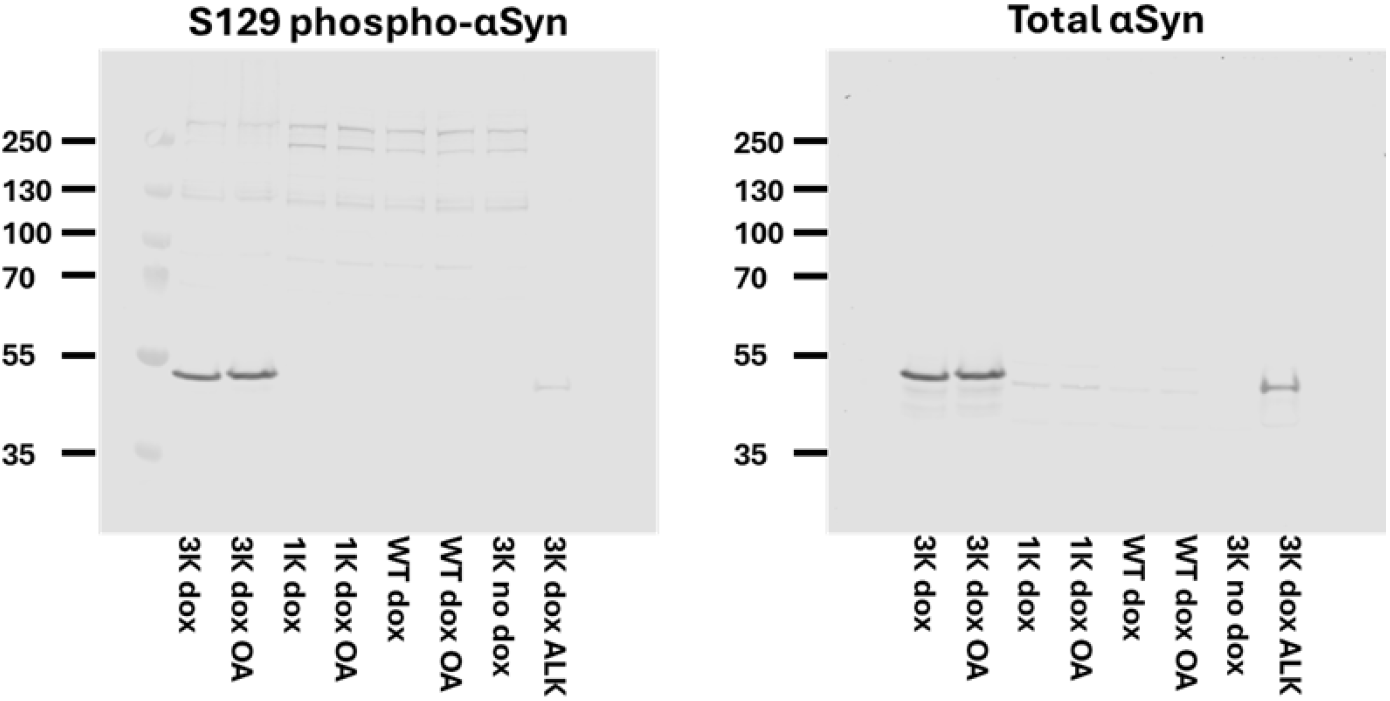
Uncropped versions of the Western blots in figure 2K.

